# Revealing Subject-Specific Temporal Patterns from Longitudinal Data

**DOI:** 10.64898/2026.02.01.703114

**Authors:** Christos Chatzis, David Horner, Rasmus Bro, Ann-Marie Malby Schoos, Morten A. Rasmussen, Evrim Acar

## Abstract

**Motivation:** Temporal multivariate data is ubiquitous in many domains, for instance, being collected over time at planned visits (every few months/years) in longitudinal cohorts, or every few minutes/hours in challenge tests. The analysis of such data often focuses on revealing the underlying temporal patterns common across subjects. However, there are subject-specific differences in temporal patterns, which hold the promise to enhance our understanding of underlying mechanisms and facilitate personalized approaches. Nevertheless, extracting subject-specific temporal patterns from longitudinal multivariate data reliably is an open challenge.

**Results:** We introduce coupled matrix factorizations (CMF) as effective tools to capture subject-specific temporal patterns focusing on two novel applications: analysis of longitudinal metabolomics data and sensitization data. Our analysis shows that CMF models reliably capture subject-specific (shape) differences in temporal patterns revealing further insights compared to the state of the art. In metabolomics, CMF models reveal differences in metabolic responses of individuals (in a postprandial meal challenge) according to anthropometric and insulin sensitivity measures. In sensitization data analysis, CMF-based methods capture differences in temporal trajectories of children according to delivery/birth mode. We demonstrate the reliability of extracted patterns using reproducibility and replicability.

**Availability:** The code is available on https://github.com/cchatzis/Revealing-Subject-specific-Temporal-Patterns-from-Longitudinal-Data. Clinical data is not publicly available due to privacy reasons. Data can be made available under a joint research collaboration by contacting COPSAC.

## 1 Introduction

Longitudinal multidimensional datasets are collected in many fields with the goal of extracting insights, capturing early risk markers of diseases or understanding various conditions, e.g., exposures in early life. For instance, blood metabolites are measured over time during meal challenge tests to understand differences in metabolic responses of individuals and how those are related to cardiometabolic diseases [Berry et al., 2020]; gut microbiome data collected over time is analyzed to understand how groups of microbes change in time and links with various conditions, e.g., delivery/birth mode [Martino et al., 2021], dietary interventions [Ma and Li, 2023]. Similarly, sensitization to allergens during childhood has been studied to investigate associations with atopic diseases [Schoos et al., 2017, Thorsen et al., 2025]. Despite differences in data characteristics, a common goal in many domains is to capture individual-specific temporal trajectories of the underlying patterns and to understand the reasons for individual differences. For instance, sensitization to certain food allergens has been shown to increase in early childhood and then decrease [Schoos et al., 2017]. The temporal trajectory, however, is not necessarily the same for everyone, and differences may reveal important insights. Therefore, a crucial question is how to reliably capture individual-specific temporal trajectories of the underlying patterns (see Figure 1).

**Fig. 1:**
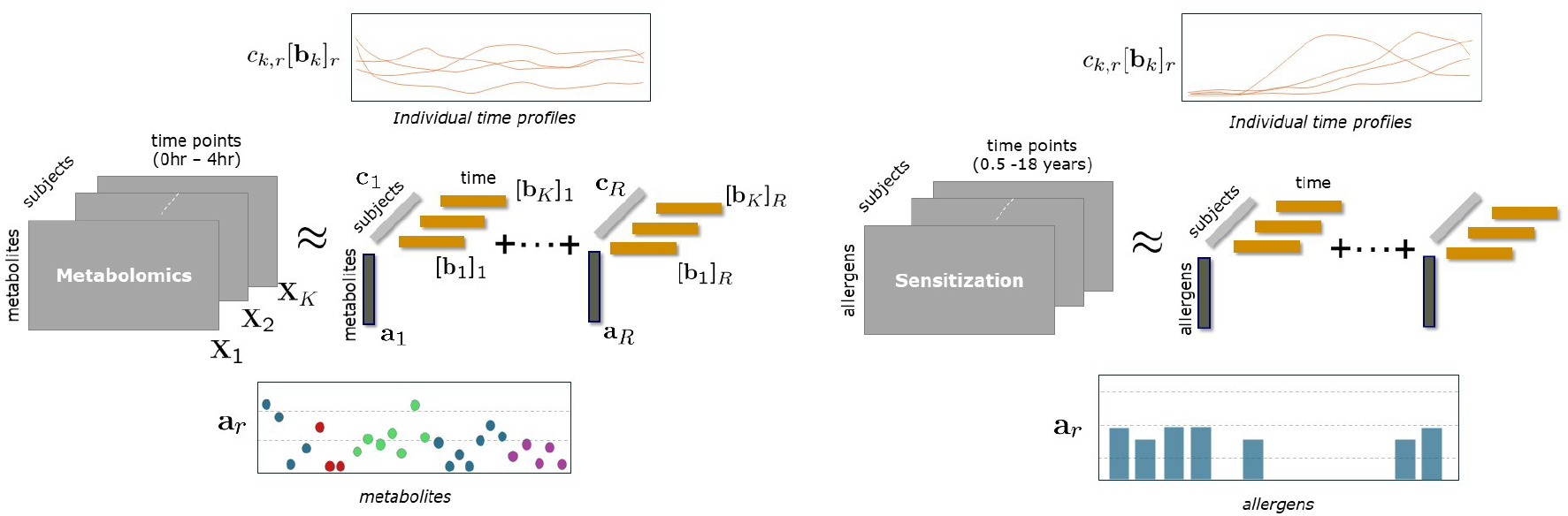
CMF models revealing subject-specific time profiles from a *metabolites* by *time* by *subjects* tensor, and an *allergens* by *time* by *subjects* tensor.

Multivariate measurements collected from multiple subjects over time form a multi-way array (in the case of aligned signals) or a collection of matrices. Traditional methods analyzing such data focus on one time point at a time, one feature at a time or summary statistics [Metwally et al., 2022, Siroux et al., 2021]. Recently, tensor decompositions such as the CANDECOMP/PARAFAC (CP) model that can reveal the underlying patterns in multi-way data have been used to analyze longitudinal multivariate data. For instance, metabolomics data in the form of *subjects* by *metabolites* by *time* tensor has been analyzed using a CP model extracting patterns in metabolite and time modes that reveal BMI (body mass index)-related differences [Yan et al., 2024]. Similarly, underlying patterns in taxa and time mode extracted from longitudinal microbiome data using CP-based approaches have revealed differences, e.g., associated with delivery/birth mode [Ma and Li, 2023, Martino et al., 2021, Shi et al., 2024, van der Ploeg et al., 2025]. CP has also been used to reveal sensitization patterns, i.e., clusters of allergens, and their temporal trajectories from longitudinal sensitization data [Schoos et al., 2017, Thorsen et al., 2025]. However, CP-based models extract common patterns across subjects and, except for a scaling difference, they do not reveal the individual variability, e.g., shape differences, in those patterns.

In this paper, our goal is to capture individual-specific temporal trajectories of the underlying patterns from longitudinal data. We demonstrate that coupled matrix factorizations (CMF) are effective tools to capture such individual differences. We focus on two novel applications: analysis of longitudinal metabolomics data and sensitization data. In metabolomics, by accounting for subject variability in temporal profiles, CMF models reveal differences in the metabolic response of individuals according to BMI and IR (insulin resistance) measures. In sensitization data analysis, CMF models reveal differences in temporal trajectories of children according to birth mode, i.e., children born by C-section show earlier sensitization to a specific group of food allergens compared to those born by natural birth. We discuss the reliability of extracted patterns in terms of reproducibility and replicability [Adali et al., 2022, Mørup et al., 2025]. Previously, PARAFAC2, which is a CMF-based approach with specific constraints and more flexible than CP, has been used to reveal subject-specific time profiles in other domains [Madsen et al., 2017, Mørup et al., 2025, Perros et al., 2019, Erdos et al., 2025]. We assess the performance of CMF-based models including PARAFAC2 and discuss our findings in comparison with CP models.

## 2 Materials and Methods

### 2.1 Datasets

We use two longitudinal datasets from the COPSAC2000 cohort, which consists of 411 subjects (with mothers with a history of asthma) [Bisgaard, 2004]. The first dataset contains metabolomics measurements of blood samples collected during a meal challenge test at the 18-year-old visit. The second dataset corresponds to allergen-specific immunoglobulin E (sIgE) measurements to allergens over the first 18 years.

#### Metabolomics Data

299 participants took part in the meal challenge test consuming a meal after overnight fasting. Blood samples were collected from participants at eight time points at 0hr (fasting state), 0.25hr, 0.5hr, 1hr, 1.5hr, 2hr, 2.5hr and 4hr after meal intake. Plasma samples were measured using NMR (nuclear magnetic resonance) spectroscopy providing a set of metabolites. We also included insulin and C-peptide hormone measurements. Details about the meal challenge test and data have previously been described [Yan et al., 2024].

Metabolomics data is arranged as a *metabolites* by *time* by *subjects* tensor (Figure 1). We removed several subjects as outliers as in previous studies [Li et al., 2024, Yan et al., 2024]. The data corresponding to males is a 161 *metabolites* by 8 *time points* by 140 *males* tensor. For females, there are two subjects with missing samples. A missing sample results in missing entries for all metabolites at a time point, which introduces a missing column in **X**_*k*_ in Figure 1. We removed those two subjects and analyzed the 161 *metabolites* by 8 *time points* by 150 *females* tensor (See Discussion for including subjects with missing samples). We analyzed males and females separately due to previously observed sex-related differences in metabolic responses [Yan et al., 2024]. Previous studies on these measurements focused on revealing subject stratifications based on fasting state data and the metabolic response in dynamic state; therefore, they analyzed fasting state-corrected data (i.e., measurements at 0hr were subtracted from other time points) [Li et al., 2024, Yan et al., 2024]. Here, we analyzed the data without fasting state correction since we are interested in individual temporal trajectories. Before the analysis, the data was scaled within metabolites mode by dividing each metabolite slice by root mean square (RMS) of that slice.

Additional data on subjects including weight, height, BMI, insulin resistance, body composition measures are available. We use these measures to study the relation between extracted patterns and phenotypes of interest (e.g., BMI-related) and to demonstrate differences in temporal trajectories of various groups, e.g., no IR, higher BMI vs. IR, higher BMI. Since there are more patterns related to BMI-related phenotypes in males than females, our experiments focus on the analysis of data from males. We discuss the results from females briefly.

Meal challenge tests generating longitudinal metabolomics data have been commonly used in nutrition and health research [Wopereis, 2022]. Individual-specific temporal trajectories of the underlying patterns extracted from the data can better characterize the difference between subject stratifications while also paving the way for personalized nutrition [Jansen et al., 2012, Wopereis, 2022]. That is what we aim to demonstrate through the metabolomics application.

#### Sensitization Data

Blood samples were collected at ages 0.5, 1.5, 4, 6, 13 and 18 years, and sIgE levels were measured against five food allergens (milk, egg, wheat flour, peanut, soybean), eight aeroallergens (birch, timothy grass, mugworth, dog, cat, horse, mold, house dust mite). Due to missing measurements for soybean and horse for everyone at the last time point, we omitted those two allergens in our analysis.

Sensitization data is arranged as a third-order tensor with modes: *allergens, time points, subjects* (Figure 1). Out of 411 children, 337 of them have a nonzero sIgE value. We included only subjects with complete information, i.e., with sIgE measurements at 6 time points for all 11 allergens. The data is a 11 *allergens* by 6 *time points* by 176 *subjects* tensor.

Sensitization is often defined as sIgE *≥* 0.35kU_A_*/*L and previous studies treated the data as binary data indicating whether there was sensitization, i.e., sIgE *≥* 0.35 or not [Schoos et al., 2017, Thorsen et al., 2025]. Rather than using a cut-off value which may have limitations [Schoos et al., 2020], we used the measured sIgE values ranging between 0 - 384.6kU_A_*/*L. The data was then preprocessed by log-transform, i.e., 𝒳_*ijk*_ = log (𝒳_*ijk*_ + 1) for the analysis not to be dominated by large values, where 𝒳_*ijk*_ is the (*i, j, k*)th entry of *allergens* by *time* by *subjects* tensor 𝒳 containing sIgE values. Before the analysis, 𝒳 was also scaled within allergens mode.

Previously, the sensitization data was analyzed using a CP model to characterize the underlying sensitization patterns, i.e., allergen groups and their temporal profiles, and to relate them to atopic diseases [Schoos et al., 2017, Thorsen et al., 2025]. These studies revealed aeroallergen and food allergen-dominant patterns and demonstrated how those patterns peaked at different ages or continued to increase over time. Those temporal profiles characterize the overall behaviour in the cohort, while individual time trajectories showing whether/when individuals grow out of allergies may be quite different. Through the analysis of sensitization data, we aim to capture such individual temporal trajectories and understand potential conditions that may lead to differences.

### 2.2 Coupled Factorizations

Let **X**_*k*_ ∈ ℝ^*I*×*J*^ correspond to longitudinal multivariate measurements from subject *k*, i.e., *I* features measured at *J* time points. For *K* subjects, matrices **X**_*k*_, *k* = 1, …, *K*, can be jointly analyzed to extract the underlying patterns using coupled factorizations. Coupled factorizations are effective approaches for capturing interpretable patterns from multi-way, multi-modal, multi-set data. They accommodate a variety of modeling assumptions, including different coupling relations [Mørup et al., 2025]. Here, we focus on CP, PARAFAC2 and CMF models, which fall under coupled factorizations, and differ in terms of coupled modes and the coupling relation between **X**_*k*_ matrices.

#### CANDECOMP/PARAFAC (CP)

Matrices **X**_*k k*=1,…,*K*_ of size *I* × *J* can be arranged a third-order tensor 𝒳 ∈ ℝ^*I*×*J*×*K*^ . Tensor factorizations have been successfully used to extract the underlying patterns from such multi-way data in many disciplines [Acar and Yener, 2009, Ballard and Kolda, 2025, Smilde et al., 2004]. One of the most commonly used tensor models is the CP model [Carroll and Chang, 1970, Harshman, 1970], which approximates the data as follows, using *R* components:

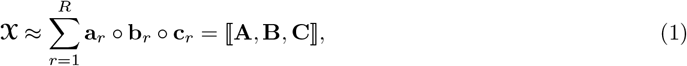

where **A** ∈ ℝ^*I*×*R*^ = [**a**_1_ **a**_2_ … **a**_*R*_], **B** ∈ ℝ^*J*×*R*^ = [**b**_1_ **b**_2_ … **b**_*R*_], and **C** ∈ ℝ^*K*×*R*^ = [**c**_1_ **c**_2_ … **c**_*R*_] correspond to factor matrices in each mode, and ◦ denotes the outer product.

In slice-wise matrix notation, the CP model corresponds to **X**_*k*_ ≈ **AD**_*k*_**B**^T^, where **D**_*k*_ ∈ ℝ^*R*×*R*^ is a diagonal matrix with *k*th row of **C** on the diagonal. If **X**_*k*_ corresponds to *allergens* by *time points* sensitization data for the *k*th subject, **a**_*r*_ captures the *r*th allergen pattern, **b**_*r*_ corresponds to its temporal profile while **c**_*r*_ shows the subject scores indicating how much *r*th pattern shows up in each subject. The CP model assumes that **X**_*k*_ matrices are coupled in both *allergens* and *time* modes, and extracts the same factor matrices **A** and **B** for all subjects, *k* = 1, …, *K*. CP is unique up to permutation and scaling ambiguities, making it possible to extract reproducible patterns from the data. Therefore, CP is often used for intepretable pattern discovery in a wide range of applications [Acar and Yener, 2009, Ballard and Kolda, 2025, Mørup et al., 2025, Smilde et al., 2004].

In our experiments, we fit CP models with nonnegativity constraints in all modes by solving the following problem:

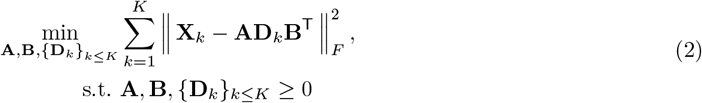

#### PARAFAC2

PARAFAC2 [Harshman, 1972] relaxes the coupling assumption in the CP model, and jointly analyzes matrices {**X**_*k*_}_*k*=1,…,*K*_ using an *R*-component model as follows:

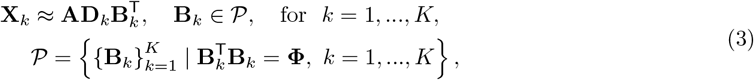

where each **X**_*k*_ is modelled using its own **B**_*k*_ ∈ ℝ^*J*×*R*^ matrix, rather than the same **B** for all **X**_*k*_s (Figure 1). Together with the cross product constraint on **B**_*k*_ matrices in (3), where **Φ** ∈ ℝ^*R*×*R*^ is the same for all *k*, the model is unique up to permutation and scaling ambiguities [Kiers et al., 1999]. For an *allergens* by *time points* sensitization matrix **X**_*k*_ for the *k*th subject, **a**_*r*_ captures the *r*th allergen pattern, [**b**_*k*_]_*r*_ corresponds to the temporal profile of that pattern for subject *k*, where [**b**_*k*_]_*r*_ denotes the *r*th column of **B**_*k*_, and **c**_*r*_ shows the subject scores for pattern *r*. As a result of its uniqueness properties which enable reproducible pattern discovery and facilitate interpretation, PARAFAC2 has been previously used in many applications. In chemometrics, it was used to separate mixtures measured using GC-MS (gas chromatography-mass spectrometry) by accounting for differences in elution profiles of chemicals across mixtures [Amigo et al., 2008]. PARAFAC2 was also used to reveal subject-specific temporal patterns in neuroimaging data analysis [Madsen et al., 2017, Mørup et al., 2025], to extract patient-specific temporal trajectories from EHR (electronic health records) [Perros et al., 2019] and to capture subject-specific temporal patterns in longitudinal microbiome data analysis [Erdos et al., 2025].

In this paper, we use PARAFAC2 models with nonnegativity constraints in all modes and solve the following problem:

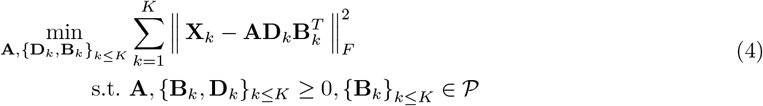

#### Coupled Matrix Factorizations (CMF)

Given matrices {**X**_*k*_} _*k*=1,…,*K*_ of size *I* × *J* coupled in one mode, e.g., the first mode, they can be jointly analyzed using coupled matrix factorizations (also known as collective matrix factorizations [Singh and Gordon, 2008]) using *R* components as follows:

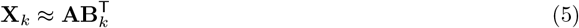

This formulation cannot uniquely recover factor matrices **A** ∈ ℝ^*I*×*R*^ and {**B**_*k*_} _*k*≤*K*_ ∈ ℝ^*J*×*R*^. However, CMF models with additional constraints have been used in various applications to find the underlying patterns, e.g., with nonnegativity constraints on all factor matrices to analyze multiple gene expression datasets [Badea, 2008], with nonnegativity and sparsity constraints on **A**, temporal smoothness on {**B**_*k*_} _*k*≤*K*_ to extract patient-specific temporal profiles in EHR analysis [Zhou et al., 2014]. In our analysis, we use CMF models with nonnegativity constraints in all modes by solving the following problem:

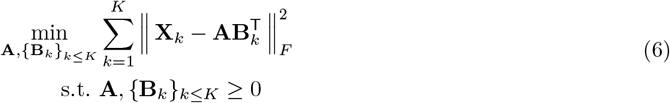

This is a coupled nonnegative matrix factorization (NMF) problem which, as we see in our reproducibility results, achieve unique results as a result of nonnegativity constraints and coupling. For **X**_*k*_ corresponding to *allergens* by *time points* sensitization data for the *k*th subject, **a**_*r*_ captures the *r*th allergen pattern while [**b**_*k*_]_*r*_ corresponds to the temporal profile of that pattern for subject *k* (as in PARAFAC2) (Figure 1). Subject scores for pattern *r* can be obtained by setting *c*_*k,r*_ = ∥ **a**_*r*_ ∥ ∥ [**b**_*k*_]_*r*_ ∥ and then normalizing **c**_*r*_. This model is less constrained than (4) since the cross-product constraint 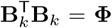 is dropped not to distort subject-specific temporal profiles due to the constraint.

### 2.3 Experimental set-up

We analyze metabolomics and sensitization data using CP, PARAFAC2 and CMF models by solving the optimization problems in (2), (4) and (6). We fit the models using different number of components (*R*), and assess their reproducibility and replicability - which is crucial for discovering patterns and to be able to interpret them [Adali et al., 2022, Mørup et al., 2025]. Reproducibility refers to the ability to uncover a unique set of patterns/components given the same data. We investigate this ability of each model by comparing the solutions obtained by different initializations reaching the same (minimum) objective function value. Replicability refers to the ability of a model to consistently recover similar patterns across different datasets. In this paper, we assess replicability by comparing the patterns extracted from different subsets of the data.

The similarity between two solutions is quantified using the following factor match scores (FMS):

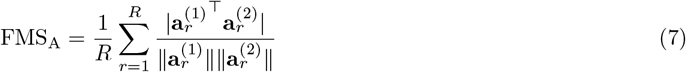

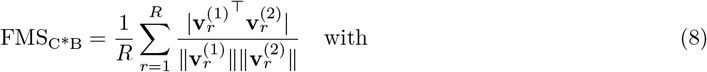

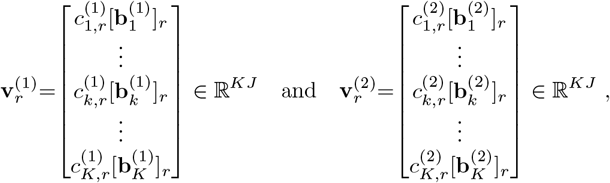

where superscripts (1) and (2) denote different factorizations (i.e., solutions), and *c*_*k,r*_[**b**_*k*_]_*r*_ denotes the scaled time profiles for the *k*th subject for the *r*th component. FMS_A_ captures the (absolute) cosine similarity between the first mode factor matrices (e.g., metabolites or allergens), FMS_C*B_ quantifies the similarity between the scaled time profiles. Both metrics are computed after matching the order of components. An FMS value of 1.0 indicates perfect match between two models.

When assessing reproducibility, we compare the solutions from different models in terms of FMS_A_ and FMS_C*B_. For replicability, we use the following procedure:

1. Randomly split the data in 10 (possibly stratified) folds
2. Form 10 subsets of the data by removing each fold from the complete dataset
3. Fit the model to each subset
4. Compare the models (in terms of FMS_A_ and FMS_C*B_) across different subsets
5. Repeat steps (1-4) 10 times.

From this procedure, a total of 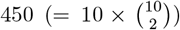 FMS_A_ and 450 FMS_C*B_ values are computed. When comparing models from different subsets of data, only the common subjects are used in FMS_C*B_ computations [Erdos et al., 2025]. We consider that models are replicable when 95% of the FMS values are above 0.90.

#### Implementation Details

For the CP model, we use `non_negative_parafac` function from TensorLy [Kos-saifi et al., 2019] using block coordinate multiplicative updates. For CMF and PARAFAC2, we use MatCoupLy [Roald, 2023], which utilizes AO-ADMM (alternating optimization alternating direction method of multipliers) [Roald et al., 2022] with Expectation Maximization (EM) to handle missing data [Chatzis et al., 2025] (missing data is only present in metabolomics, i.e., 0.07% of the entries in males and 0.08% of the entries in females).

For each experiment, 50 random initializations were used, and the best run was chosen as the one reaching the minimum function value after discarding runs that reached the maximum number of iterations (set to 15000). For AO-ADMM, all runs with feasibility gaps larger than 10^−5^ at termination were discarded. The algorithms stopped due to either small absolute (10^−6^) or relative (10^−8^) change in the total loss.

## 3 Results

### 3.1 Longitudinal Metabolomics Data Analysis

Figure 2 shows the components extracted using a 6-component CMF model of tensor 𝒳 by solving (6), where **X**_*k*_s correspond to 161 metabolites × 8 time points matrices, and *K* = 140 subjects (See Reproducibility and Replicability for the selection of number of components). For each component *r*, the metabolite pattern (**a**_*r*_) and subject-specific time profiles scaled by subject scores (*c*_*k,r*_[**b**_*k*_]_*r*_) are shown. Metabolites are colored according to lipoprotein classes, i.e., HDL (high density lipoprotein), IDL (intermediate density lipoprotein), LDL (low density lipoprotein), and VLDL (very low density lipoprotein). Glycolysis-related metabolites, ketone bodies, insulin, and c-peptide are also marked. The last column shows the scaled subject-specific time profiles colored according to groups considering both BMI and IR, where BMI and IR groups are defined as: lower BMI: BMI < 25, higher BMI: BMI ≥ 25; NoIR: HOMA-IR ≤ 2.91, IR: HOMA-IR > 2.91 [Silva et al., 2022].

**Fig. 2:**
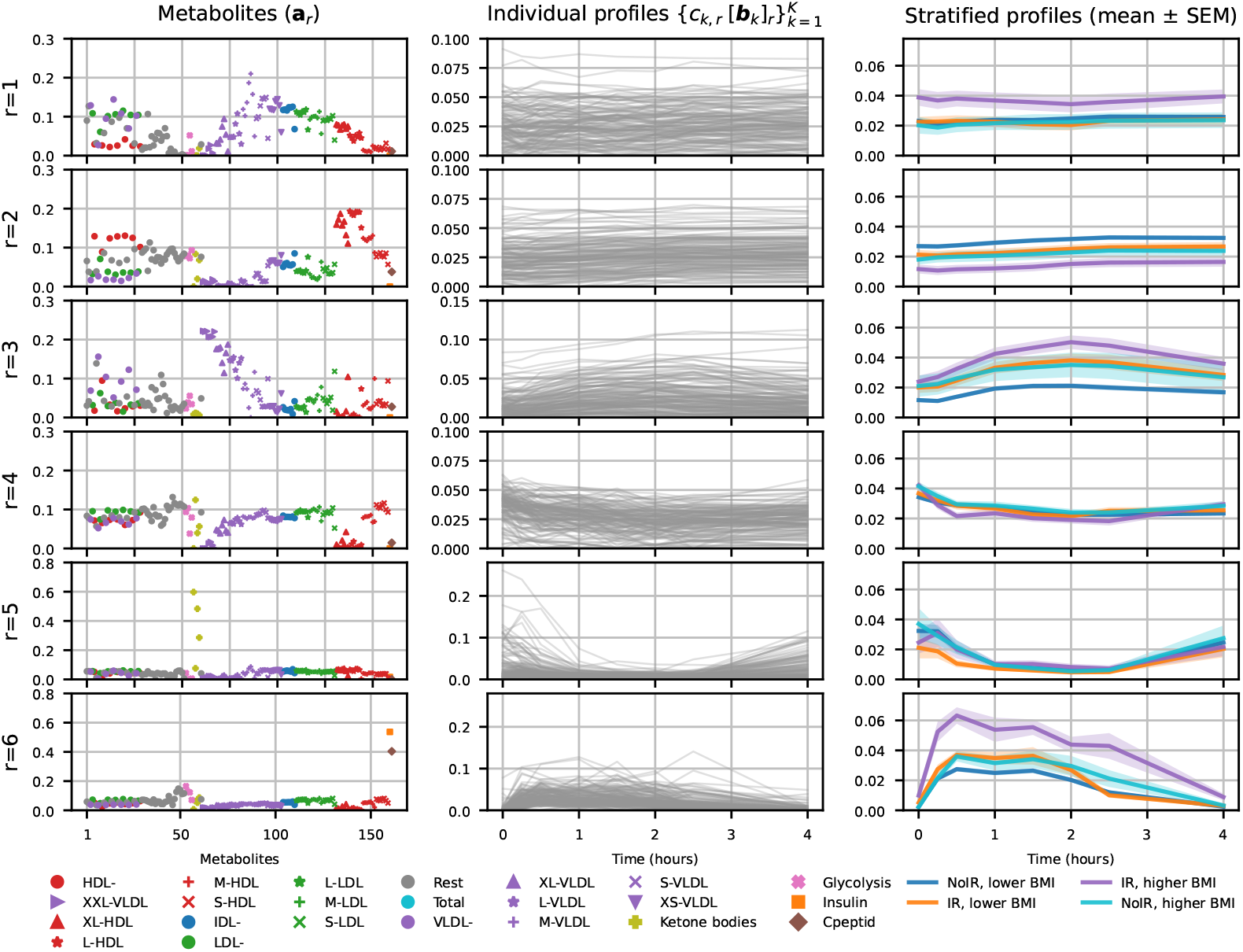
Metabolomics. Components of a 6-component CMF model (with nonnegativity constraints in all modes) of the metabolomics data from males. **a**_*r*_ denotes the pattern in the metabolites mode, where metabolites are colored by lipoprotein classes. Different shapes are used for lipoprotein subclasses. Subject-specific time profiles scaled by the corresponding subject scores, i.e., *c*_*k,r*_[**b**_*k*_]_*r*_, for each component are shown in the middle column. The last column shows scaled subject-specific time profiles colored according to four BMI/IR groups. On the y-axis, the label *r* = 1 denotes component 1, *r* = 2 denotes component 2 etc.

Out of six components, subject scores (**c**_*r*_) of four components (comp. 1, 2, 3 and 6) show statistically significant difference in terms of BMI groups. Component 1 mainly captures the response of smaller size (XS, S, M) VLDL, IDL and LDL while component 3 models larger size (XXL, XL, L) VLDL. Subject-specific time profiles show the difference in the response of each subject in terms of these metabolites. In the third column, we observe that both BMI and IR play a role in these differences. Component 2 models HDL, and at the group level, we observe the reverse ordering of BMI/IR subgroups - with HDL being negatively associated with the (IR, higher BMI) group. Component 6 captures the response of insulin, c-peptide and, also picks up glycolysis-related metabolites (glucose, lactate, pyruvate) and some aminoacids (isoleucine, leucine). The second column shows the individual differences in the time profiles, and the last column shows how the response peaks and decreases for different groups. Among the four components related to BMI, component 6 achieves the highest correlation. Figure 3 shows the correlations between subject scores (**c**_6_) and BMI (and other variables of interest). We have not observed any association between these variables and comp. 4 and 5. In females, similar components as in Figure 2 are revealed using a 6-component CMF model. Despite similar components (modelling lipoproteins), only the component modelling insulin, c-peptide and glycolysis-related metabolites has good correlations with BMI (and other variables) supporting previously-reported sex-differences [Yan et al., 2024] (Due to space limitations, we do not include the results from females here and provide the models in Figure S.5, S.6 and S.7 in the supplementary material.)

**Fig. 3:**
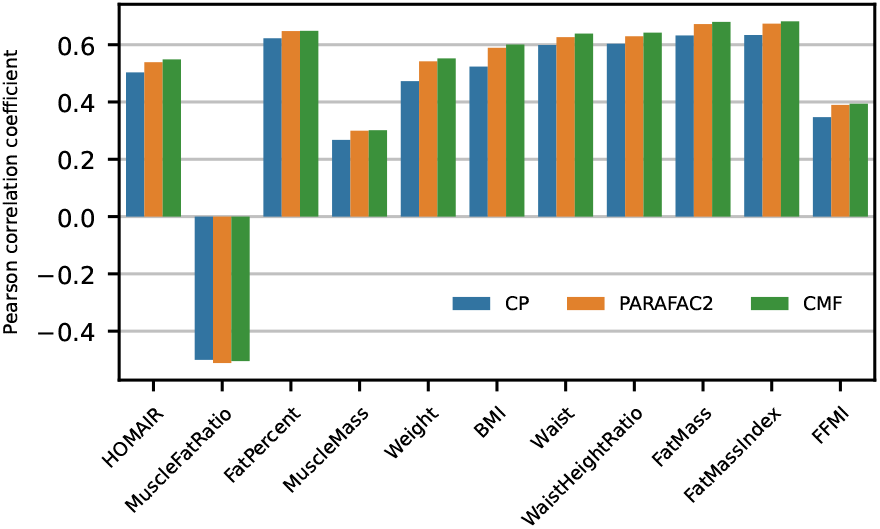
Correlation between variables of interest and (male) subject scores (**c**_*r*_) for the component modelling insulin, c-peptide and glycolysis-related metabolites. HOMA-IR: Homeostatic Model Assessment for Insulin Resistance; MuscleFatRatio denotes muscle to fat ratio; FatPercent denotes body fat percentage; MuscleMass indicates the amount of muscle in the body; Waist denotes the waist circumference; WaistHeightRatio is the waist measurement divided by height; FatMass denotes the amount of body fat; FatMassIndex is computed as FatMass/height^2^, and FFMI indicates the fat free mass index. See [Yan et al., 2024] for more details.

We compare the CMF model with a 6-component CP and a 6-component PARAFAC2 model of tensor 𝒳 (See Reproducibility and Replicability for component number selection). In the metabolites mode, both CP and PARAFAC2 capture patterns similar to those captured by CMF. Figure 4 shows the component modelling insulin, c-peptide, and glycolysis-related metabolites captured by CP and PARAFAC2 (For all components, see Figure S.2 and S.3 in the supplementary material). In terms of time profiles, CP extracts the same time profile for every subject; therefore, the response has the same shape for everyone and is only scaled by subject scores. The model fails to capture shape differences between individuals and between BMI/IR groups. PARAFAC2, on the other hand, extracts subject-specific profiles and captures shape differences between BMI/IR groups (similar to comp. 6 in the CMF model). Those shape differences may reveal biologically-relevant insights. A healthy insulin curve after a meal is normally biphasic, showing rapid insulin secretion followed by slower insulin release. This biphasic response is also observed for glucose and c-peptide. When the biphasic response turns into a monophasic response, this has been associated with insulin sensitivity [Kim et al., 2016]. We may be observing these differences in the temporal patterns when we compare CP vs. CMF/PARAFAC2. However, these findings should be considered hypothesis-generating and warrant further investigation in independent cohorts. Figure 3 shows the correlations between subject scores for the component modelling primarily insulin and c-peptide, and variables of interest. All methods achieve similar correlations indicating that subject scores from different models are similar and not enough to capture such differences in temporal trajectories.

**Fig. 4:**
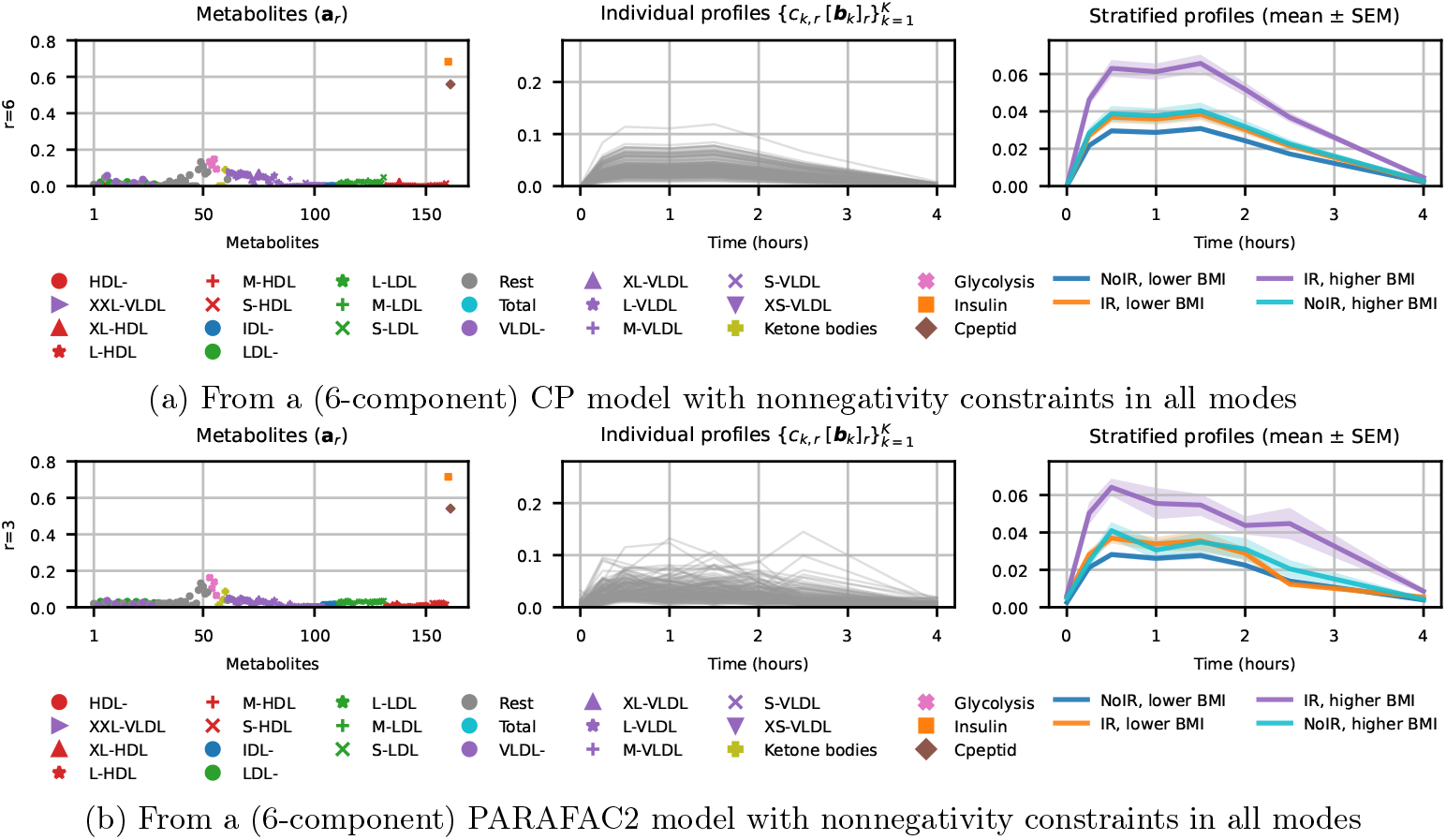
The component modelling insulin, c-peptide and glycolysis-related metabolites extracted using different models. **a**_*r*_ denotes the pattern in the metabolites mode. Subject-specific time profiles scaled by the corresponding subject scores, i.e., *c*_*k,r*_[**b**_*k*_]_*r*_, are shown in the middle column. Scaled subject-specific time profiles colored according to BMI/IR groups are shown in the last column. Note that due to permutation ambiguity, components in different models may have different orders. Therefore, this component corresponds to component 6 (marked with *r* = 6 on y-axis label) for CP while it is component 3 (*r* = 3) in PARAFAC2.

Previous studies analyzed these measurements using CP focusing on fasting state - corrected data [Yan et al., 2024], and its joint analysis with fasting data [Li et al., 2024]. They revealed BMI-related components dominated by lipoproteins as in comp. 1, 2 and 3 in Figure 2. However, the component modelling insulin, c-peptide and glycolysis-related metabolites was not captured achieving lower correlations with variables of interest. In this paper, we capture the response of these important metabolites, and also, via PARAFAC2 and CMF, the subject-specific time profiles.

#### Reproducibility and Replicability

Figure 5 (a) shows the reproducibility of patterns extracted by CP, PARAFAC2 and CMF models using different number of components. With FMS_A_ and FMS_C*B_ close to 1, all models with 2-8 components produce unique results and are reproducible. Figure 5 (b) shows the replicability of extracted patterns across different subsets of the data obtained via stratified sampling based on BMI groups. Except for PARAFAC2 with 8 components, all models can be considered replicable. We choose *R* = 6 for all models based on replicability and interpretation of components. Up to 5 components, insulin/c-peptide component cannot be captured. With *R* = 6, a cleaner insulin/c-peptide component is captured, compared to *R* = 5. However, for *R* = 7, we see components splitting into multiple components.

**Fig. 5:**
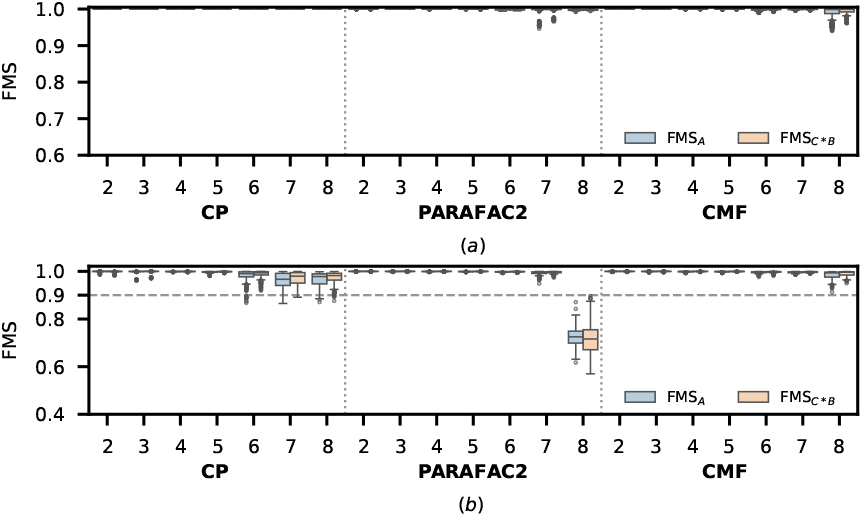
(a) Reproducibility, and (b) Replicability of different models of the metabolomics data using different number of components (*R*).

### 3.2 Longitudinal Sensitization Data Analysis

Figure 6 shows the components of a 5-component CMF model of tensor 𝒳, where **X**_*k*_ is a 11 allergens by 6 time points matrix, for *k* = 1, .., *K*, with *K* = 176 subjects (See Reproducibility and Replicability for component number selection). For each component *r*, the allergen pattern (**a**_*r*_), scaled subject-specific time profiles (*c*_*k,r*_[**b**_*k*_]_*r*_) and the mean profile of scaled subject-specific time profiles are shown. In the last column, scaled subject-specific time profiles are colored according to the delivery/birth mode, i.e., natural birth (117 subjects), C-section (35 subjects) and vacuum extraction (24 subjects).

**Fig. 6:**
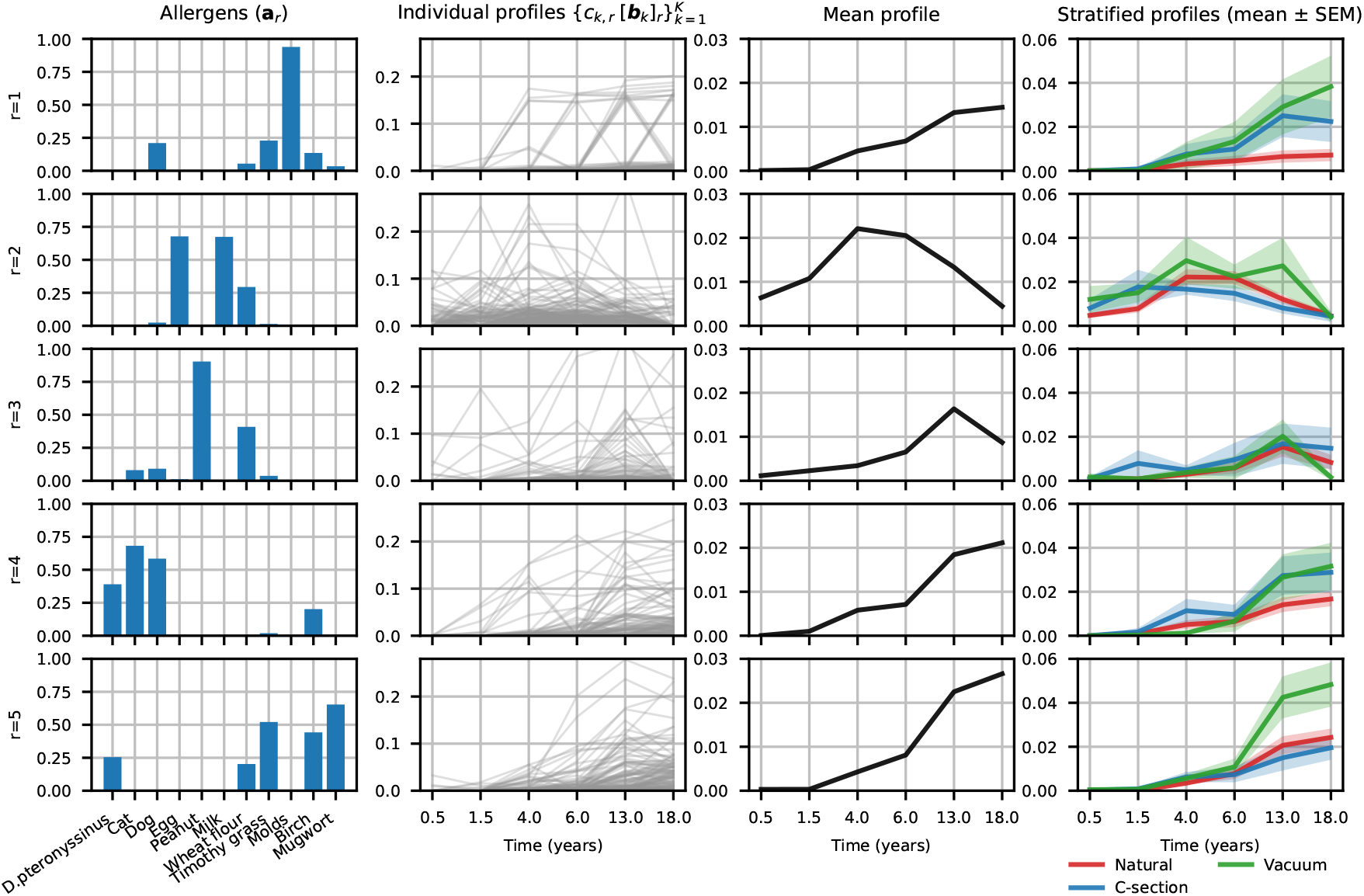
Sensitization. Components of a 5-component CMF model (with nonnegativity constraints in all modes) of the sensitization data. **a**_*r*_ denotes the pattern in the allergens mode. Subject-specific time profiles scaled by the corresponding subject scores, i.e., *c*_*k,r*_[**b**_*k*_]_*r*_, are shown in the middle column. Mean of scaled subject-specific profiles are plotted in the third column. The last column shows mean (and standard error of mean) of scaled subject-specific time profiles colored according to delivery/birth mode groups. D.pteronyssinus denotes house dust mite.

Among the five components, two of them mainly model food allergens (**a**_2_, **a**_3_), two of them model aeroallergens (**a**_1_, **a**_4_), and one of them (**a**_5_) contains allergens from both groups. Component 2 models, in particular, the sensitization to egg, milk and wheat flour. In the second column, we observe that individuals differ in terms of their sensitization to these allergens over time. In the third column, the average profile shows that, in average, the sensitization peaks around 4 years old and then decreases. In the last column, time profiles colored according to birth mode show different peaks at different ages for different groups - revealing group differences. Similarly, component 3 mainly models the sensitization to peanut (and also wheat flour), where in average sensitization peaks around 13 years old but also showing group differences in the last column. Component 1 primarily models mold (also dog, grass, and birch) with increasing sensitization over time in average. Component 4 models house dust mite, dog, cat, and, to some extent, birch sensitization also increasing over time. Finally, the last component models both mugworth, birch, grass, house dust mite and also wheat flour showing increasing sensitization over time.

We compare the CMF model with a 5-component CP and 5-component PARAFAC2 model. All models reveal similar components in the allergens mode; however, they differ in terms of temporal trajectories. Figure 7 shows the component modelling the sensitization to egg, milk and wheat flour extracted by each model (For all components, see Figure S.8 and S.9 in the supplementary material). The CP model extracts the same temporal profile from each subject, only differing up to scaling. However, both CMF and PARAFAC2 can reveal shape differences in temporal profiles indicating subjects born by C-section showing earlier sensitization to these allergens than those born by natural birth.

**Fig. 7:**
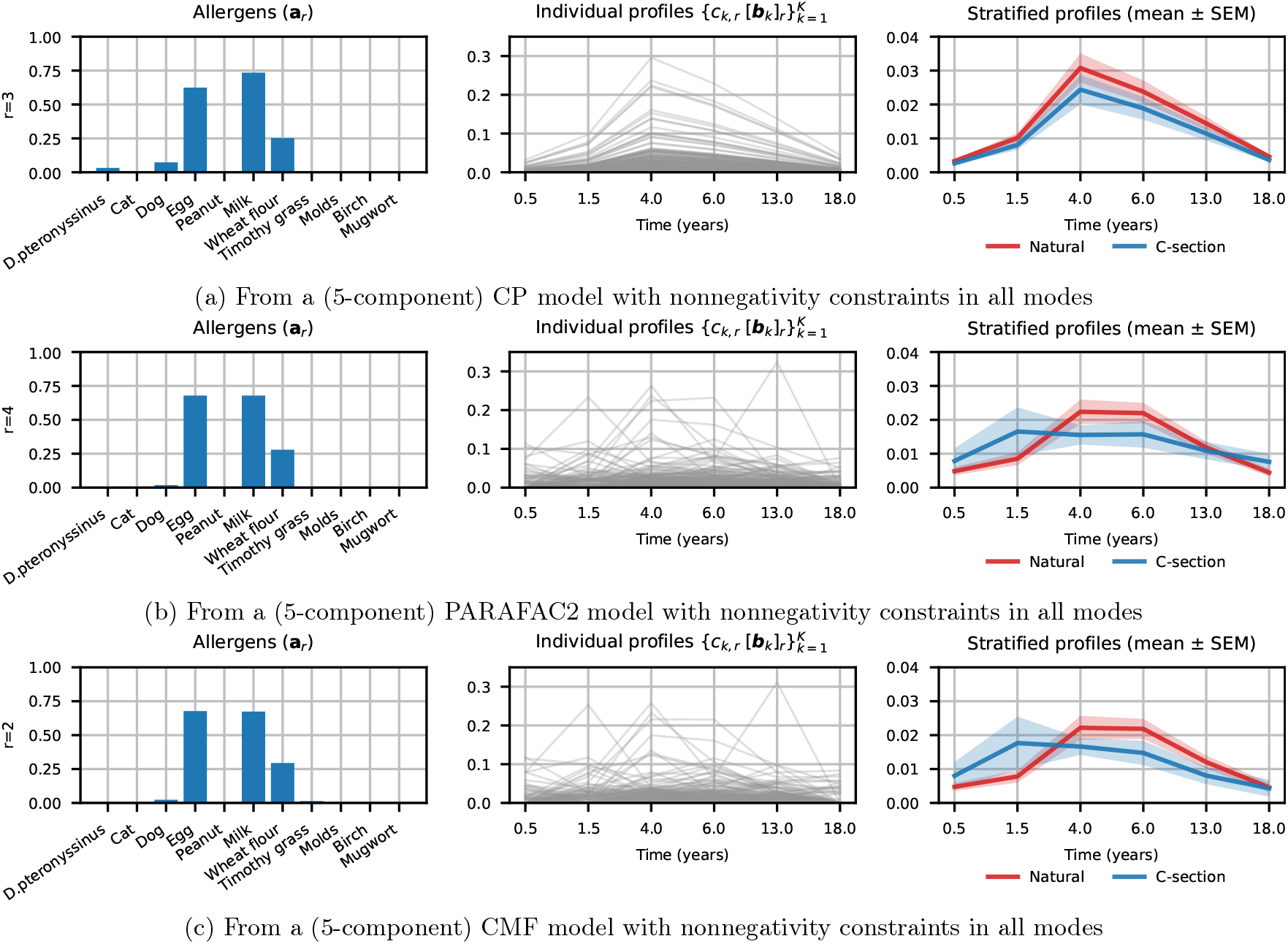
The food-allergen (i.e., egg, milk, and wheat flour) dominated component extracted using different models. **a**_*r*_ denotes the pattern in the allergens mode. Scaled subject-specific time profiles are shown in the middle column. The last column shows mean (and standard error of mean) patterns of scaled subject-specific time profiles colored according to delivery/birth mode. We only plot natural birth and C-section groups for clarity in the last column.

Compared to previous studies [Thorsen et al., 2025], CP reveals similar patterns such as components that model food allergens and their average time profiles, and components that model aeroallergens and their average time profile increasing over time. Due to sparsity constraints and the higher number of components, sparser components have previously been observed splitting some components that model aeroallergens [Thorsen et al., 2025]. However, the overall interpretation remains the same, confirming previous findings. In addition, in this paper, through the CMF and PARAFAC2 models, we reveal subject-specific sensitization trajectories showing that individuals have different temporal trajectories of sensitization and that the difference may be, for instance, related to the delivery mode. Early-life risk factors for food allergies have previously been studied showing positive associations between cesarean delivery and food allergies, with an increased risk in the c-section group already when infants were less than one year old [Mitselou et al., 2018], and also in infants with allergic predisposition [Eggesbø et al., 2003]. Our results are in line with these findings, and in addition, we provide more insights such as the allergen pattern with the specific food allergens as well as complete sensitization trajectories.

#### Reproducibility and Replicability

Figure 8 (a) shows that we consistently obtain the same solution for all models for different number of components, except for PARAFAC2 when *R* = 6 (In this case, no runs resulted in function values close to each other; therefore, 6-component PARAFAC2 was not included in the analysis). Figure 8 (b) shows the replicability of patterns, using stratified subsets according to delivery mode. We observe that CMF and PARAFAC2 are more replicable than CP. We choose *R* = 5 for CMF and PARAFAC2 based on both replicability and interpretation of the components. CMF is replicable at *R* = 5 (for higher number of components, allergen components are splitted), and PARAFAC2 is replicable only up to *R* = 5. For CP, with *R* = 5, we get allergen components clustering allergens meaningfully as in other models; therefore, for comparisons we used the 5-component CP model but note that the model is not as replicable as other models.

**Fig. 8:**
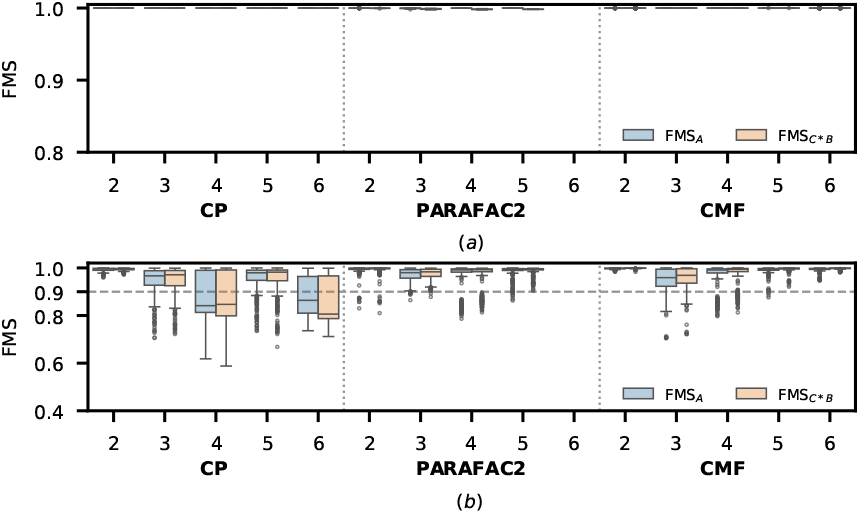
(a) Reproducibility, and (b) Replicability of different models of the sensitization data using different number of components (*R*).

## 4 Discussion

We observe that CMF and PARAFAC2 reveal similar components in terms of both features (allergens or metabolites) and subject-specific time profiles in different applications. However, while results look similar at the group level in Figure 2 vs. Figure 4 for metabolomics data analysis, and in Figure 7 for sensitization analysis, there are differences in scaled subject-specific time profiles. PARAFAC2 has an additional constraint, which comes with assumptions about orientations of components with respect to each other. When scaled subject-specific time profiles for each subject are compared, while the models (CMF vs. PARAFAC2) agree well for the components we have highlighted, there are some components where the profiles for subjects do not match well (See Figure S.1 and S.10 in the supplementary material). We consider CMF with nonnegativity constraints a better choice not to distort the time profiles with additional constraints. However, if the data is not nonnegative (due to centering or transformations [Erdos et al., 2025]), then one should consider PARAFAC2.

A common problem in longitudinal data analysis is missing samples. If subject *k* misses a sample at a time point, this corresponds to a missing column in **X**_*k*_. In the metabolomics data, there are a few such subjects. In sensitization data, out of 411 subjects, there are 77 subjects missing one sample (i.e., one visit), 29, 28, 11, and 4 subjects missing two, three, four, and five samples, respectively. Missing columns are particularly challenging for CMF-based approaches including PARAFAC2 because such missing data results in non-unique models. In this study, we excluded anyone with missing samples for the patterns/findings not to be affected by missing data. We plan to address this issue via additional constraints such as temporal smoothness in future work.

In both applications, we observe that individual temporal profiles vary widely. To understand the main sources of variation among individual temporal profiles, we have looked into various groups of interest such as BMI/IR groups in metabolomics, and delivery mode in sensitization data analysis. However, a more systematic approach is needed and we plan to explore that via joint analysis of longitudinal data and other relevant data (e.g., variables of interest, other omics data) through coupled matrix and tensor factorizations [Schenker et al., 2025].

Subject-specific temporal profiles of the underlying patterns from longitudinal data are meaningful only when they can be extracted reliably. We have demonstrated both reproducibility and replicability of those patterns in our analysis. However, our replicability analysis has been limited since it relies on the replicability of patterns across subsets of the data. In other domains, it is possible to collect replicates from the same individuals, and assess replicability using those replicates such as in functional neuroimaging [Mørup et al., 2025]. However, this approach remains a challenging issue in domains where measurements are expensive.

In conclusion, this paper demonstrates that CMF-based methods are effective approaches to reliably capture subject-specific temporal patterns via two novel applications from metabolomics and sensitization data analysis. Our results show that those individual (shape) differences may reveal important insights, e.g., links with earlier exposures such as the delivery type or various phenotypes and diseases. We have assessed the reliability of the patterns considering both reproducibility and replicability. Validation of such insights using independent cohorts remains as future work.

## Acknowledgments

We would like to thank Suzan Wopereis and Balazs Erdos for helpful discussions. We also thank the children and families of the COPSAC2000 cohort and the clinical team at COPSAC.

