## Supplementary Figures for "Revealing Subject-Specific Temporal Patterns from Longitudinal Data"

### Supplementary Material for "Revealing Subject-Specific Temporal Patterns from Longitudinal Data"

#### Metabolomics Data Analysis

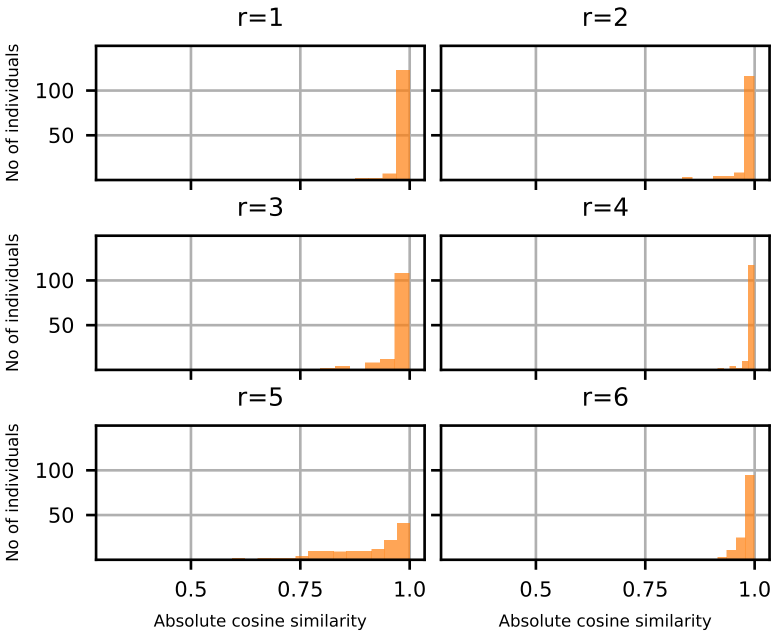


Figure S.1: Males. Comparison of subject-specific time profiles ($c_{k,r}[\mathbf{b}_k]_r$) captured by CMF and PARAFAC2 for each component for the metabolomics data. Each histogram contains 140 data points, one for each subject. The order is adjusted (following the order in the CMF model) so that matching components from different models are compared.

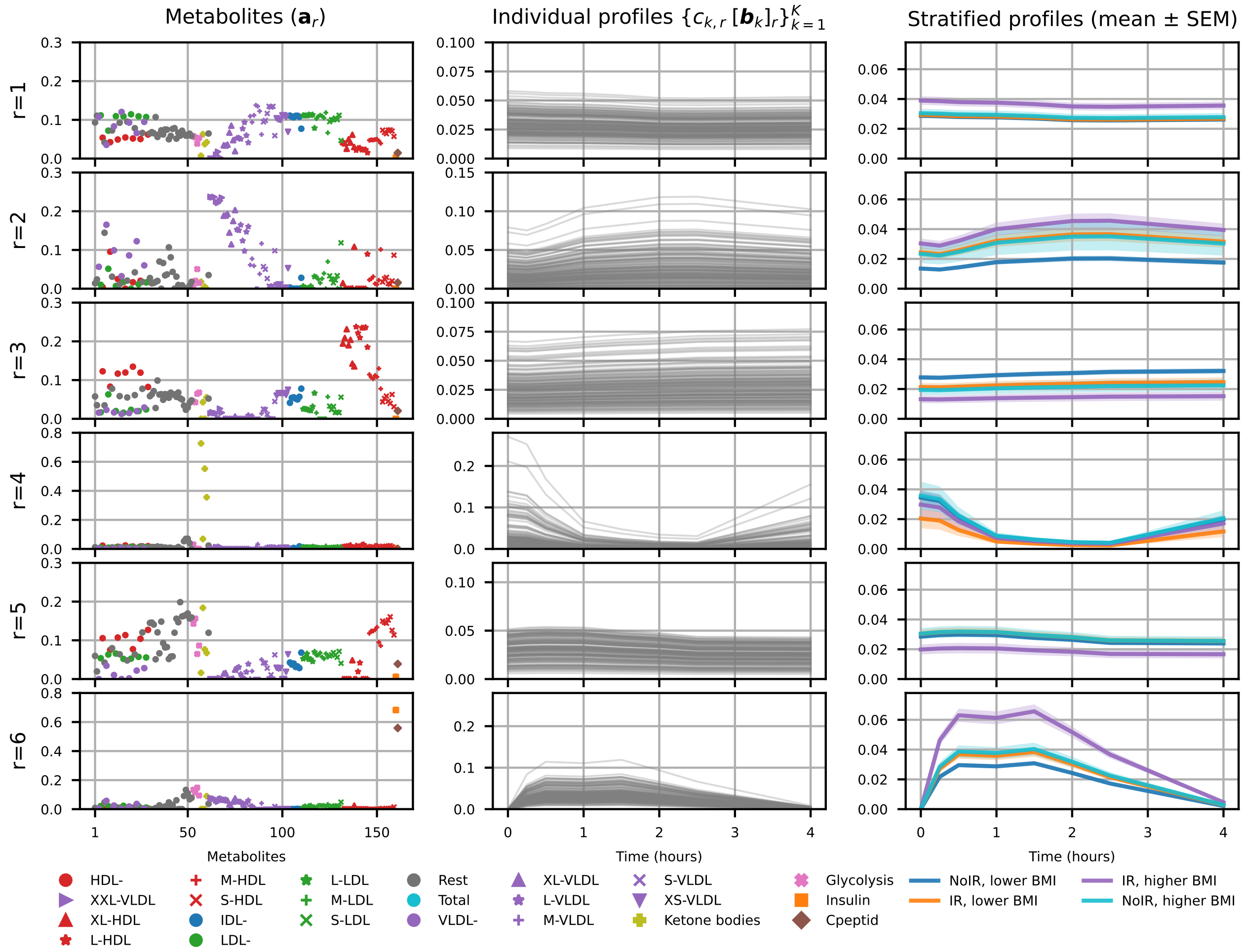


Figure S.2: Males. Components of a 6-component CP model (with nonnegativity constraints in all modes). $\mathbf{a}_r$ denotes the pattern in the metabolites mode, where metabolites are colored by lipoprotein classes. Different shapes are used for lipoprotein subclasses. Subject-specific time profiles scaled by the corresponding subject scores, i.e., $c_{k,r}[\mathbf{b}_k]_r$ for each component are shown in the middle column. The last column shows scaled subject-specific time profiles colored according to four BMI/IR groups.

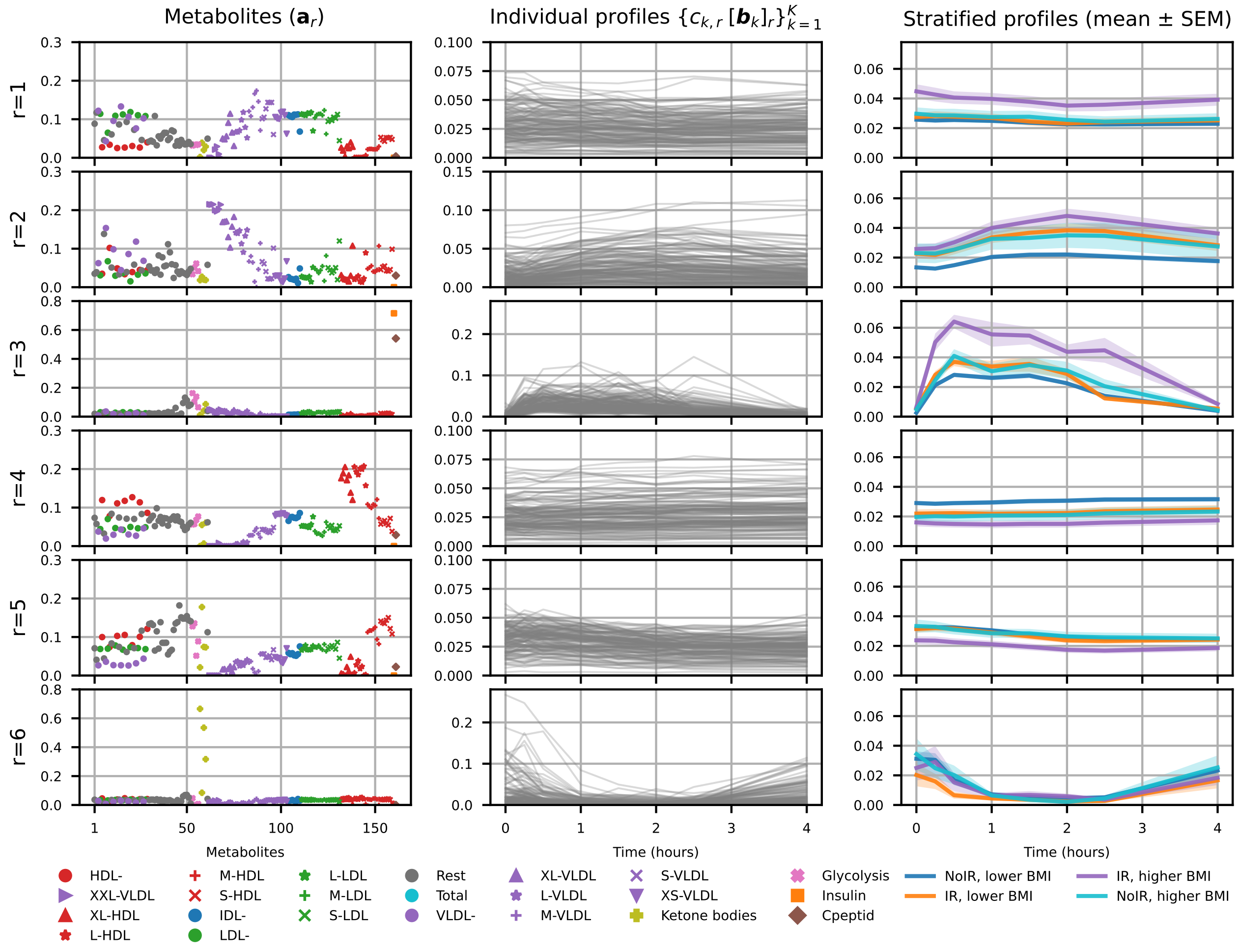


Figure S.3: Males. Components of a 6-component PARAFAC2 model (with nonnegativity constraints in all modes). $\mathbf{a}_r$ denotes the pattern in the metabolites mode, where metabolites are colored by lipoprotein classes. Different shapes are used for lipoprotein subclasses. Subject-specific time profiles scaled by the corresponding subject scores, i.e., $c_{k,r}[\mathbf{b}_k]_r$ for each component are shown in the middle column. The last column shows scaled subject-specific time profiles colored according to four BMI/IR groups.

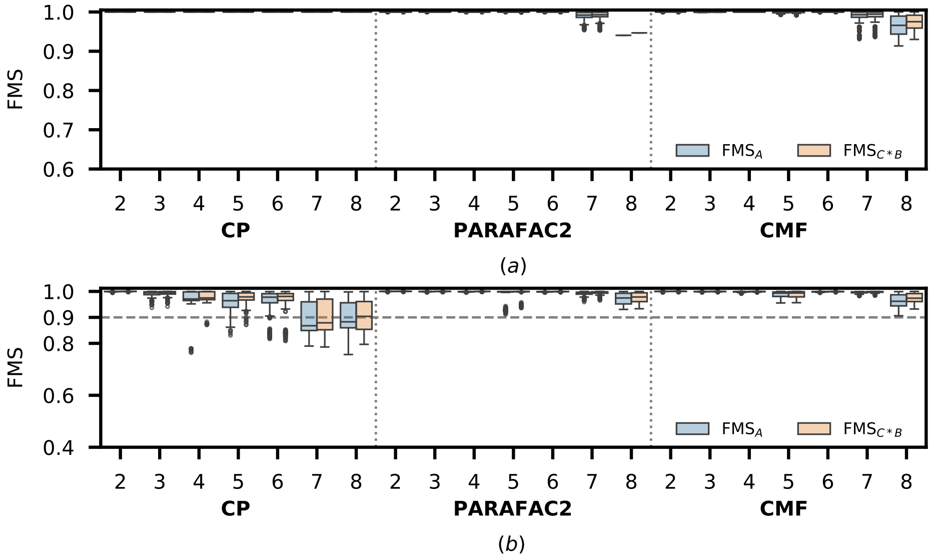


Figure S.4: Females. (a) Reproducibility, and (b) Replicability of different models of the metabolomics data using different number of components ($R$).

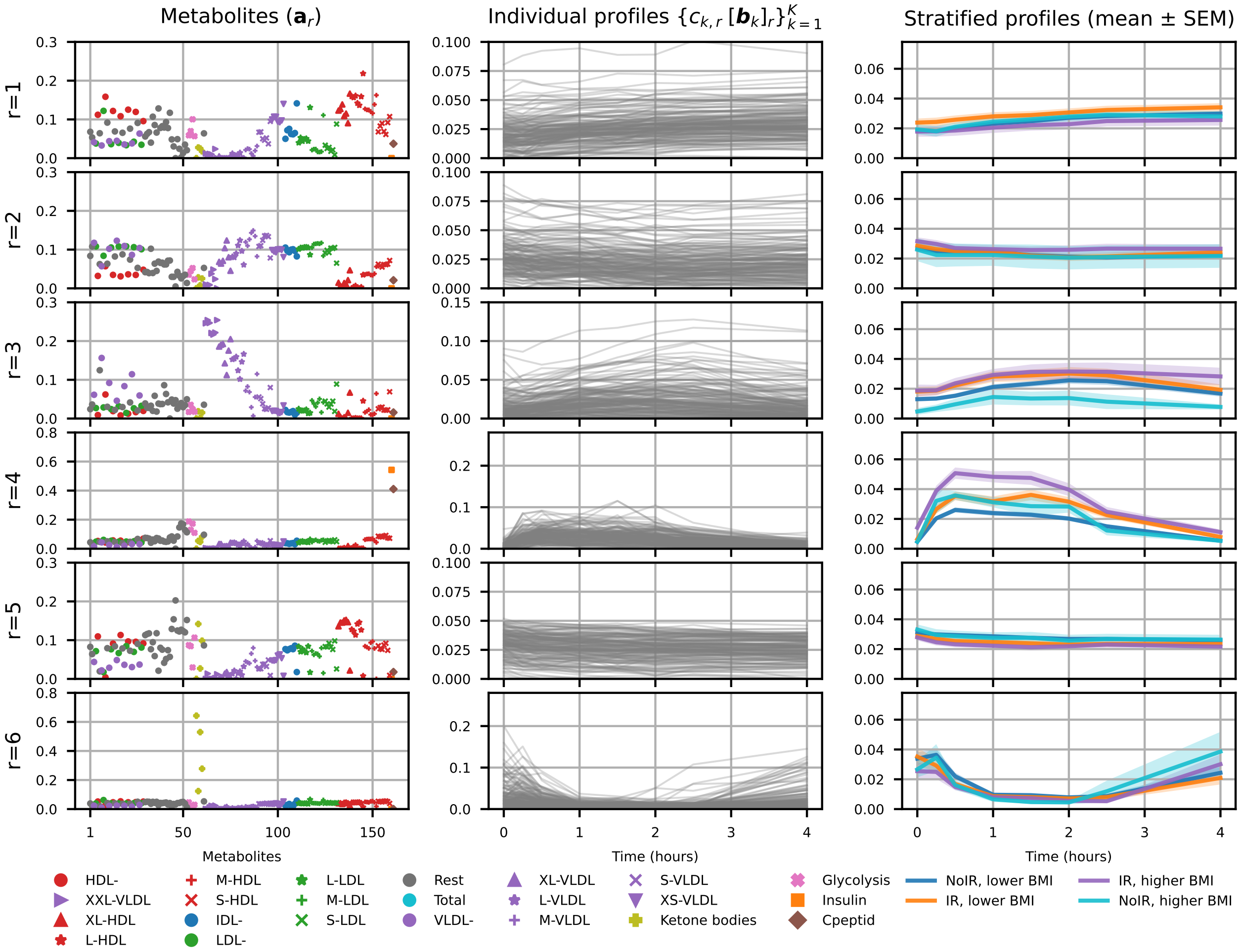


Figure S.5: Females. Components of a 6-component CMF model (with nonnegativity constraints in all modes). $\mathbf{a}_r$ denotes the pattern in the metabolites mode, where metabolites are colored by lipoprotein classes. Different shapes are used for lipoprotein subclasses. Subject-specific time profiles scaled by the corresponding subject scores, i.e., $c_{k,r}[\mathbf{b}_k]_r$ for each component are shown in the middle column. The last column shows scaled subject-specific time profiles colored according to four BMI/IR groups.

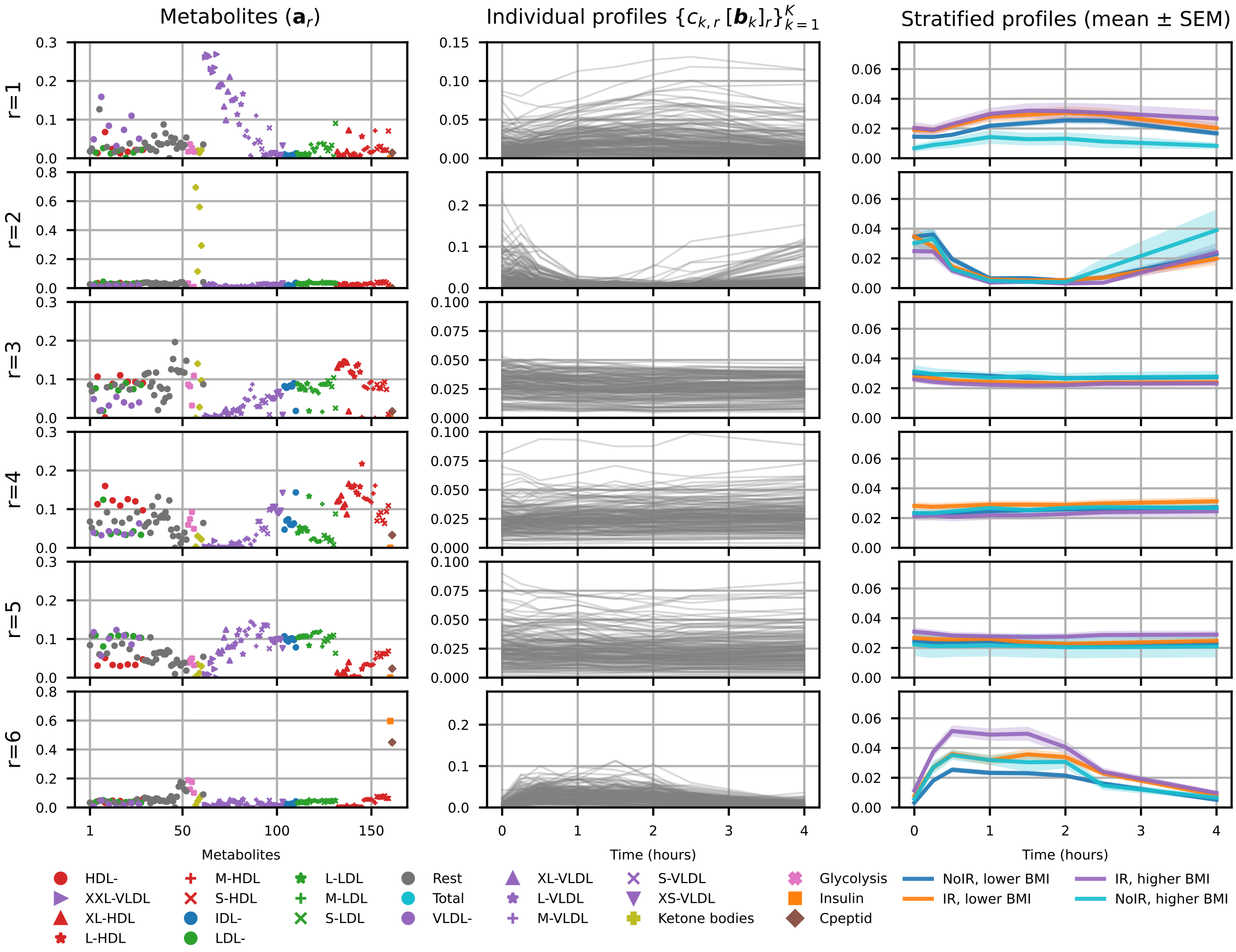


Figure S.6: Females. Components of a 6-component PARAFAC2 model (with nonnegativity constraints in all modes). $\mathbf{a}_r$ denotes the pattern in the metabolites mode, where metabolites are colored by lipoprotein classes. Different shapes are used for lipoprotein subclasses. Subject-specific time profiles scaled by the corresponding subject scores, i.e., $c_{k,r}[\mathbf{b}_k]_r$ for each component are shown in the middle column. The last column shows scaled subject-specific time profiles colored according to four BMI/IR groups.

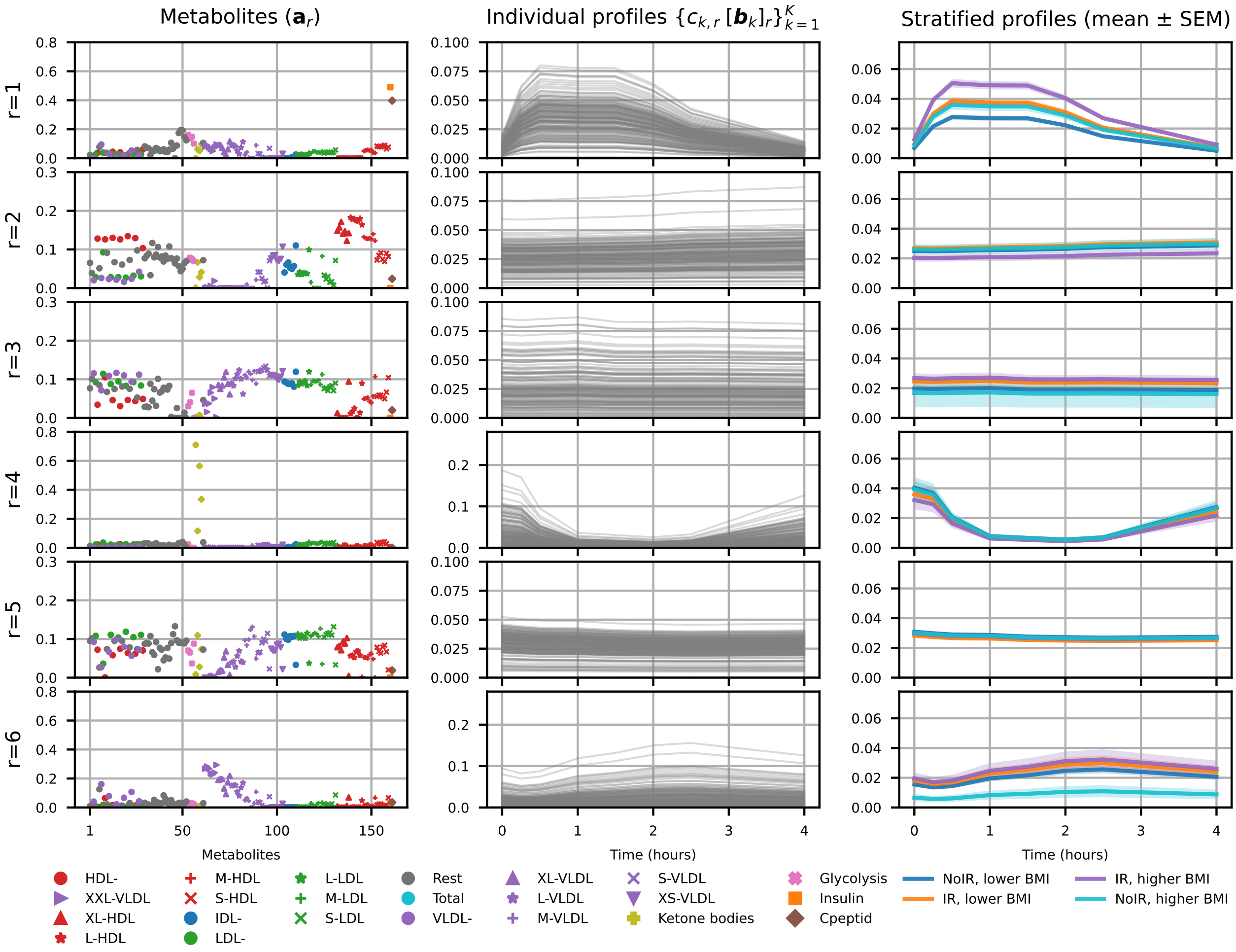


Figure S.7: Females. Components of a 6-component CP model (with nonnegativity constraints in all modes). $\mathbf{a}_r$ denotes the pattern in the metabolites mode, where metabolites are colored by lipoprotein classes. Different shapes are used for lipoprotein subclasses. Subject-specific time profiles scaled by the corresponding subject scores, i.e., $c_{k,r}[\mathbf{b}_k]_r$ for each component are shown in the middle column. The last column shows scaled subject-specific time profiles colored according to four BMI/IR groups.

### Sensitization Data Analysis

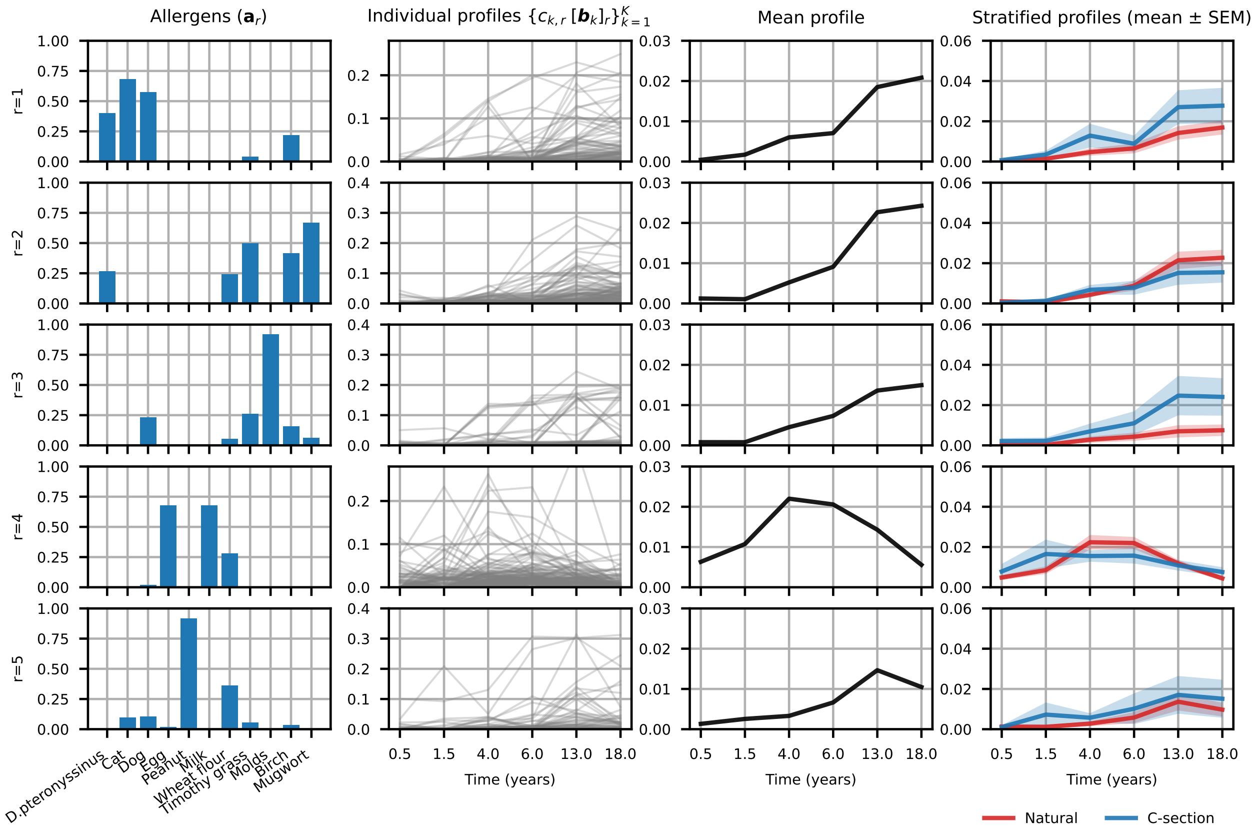


Figure S.8: Sensitization. Components of a 5-component PARAFAC2 model (with nonnegativity constraints in all modes). $\mathbf{a}_r$ denotes the pattern in the allergens mode. Subject-specific time profiles scaled by the corresponding subject scores, i.e., $c_{k,r}[\mathbf{b}_k]_r$ are shown in the middle column. Mean of scaled subject-specific profiles are plotted in the third column while the last column shows mean (and standard error of mean) patterns of scaled subject-specific time profiles colored according to delivery/birth mode groups.

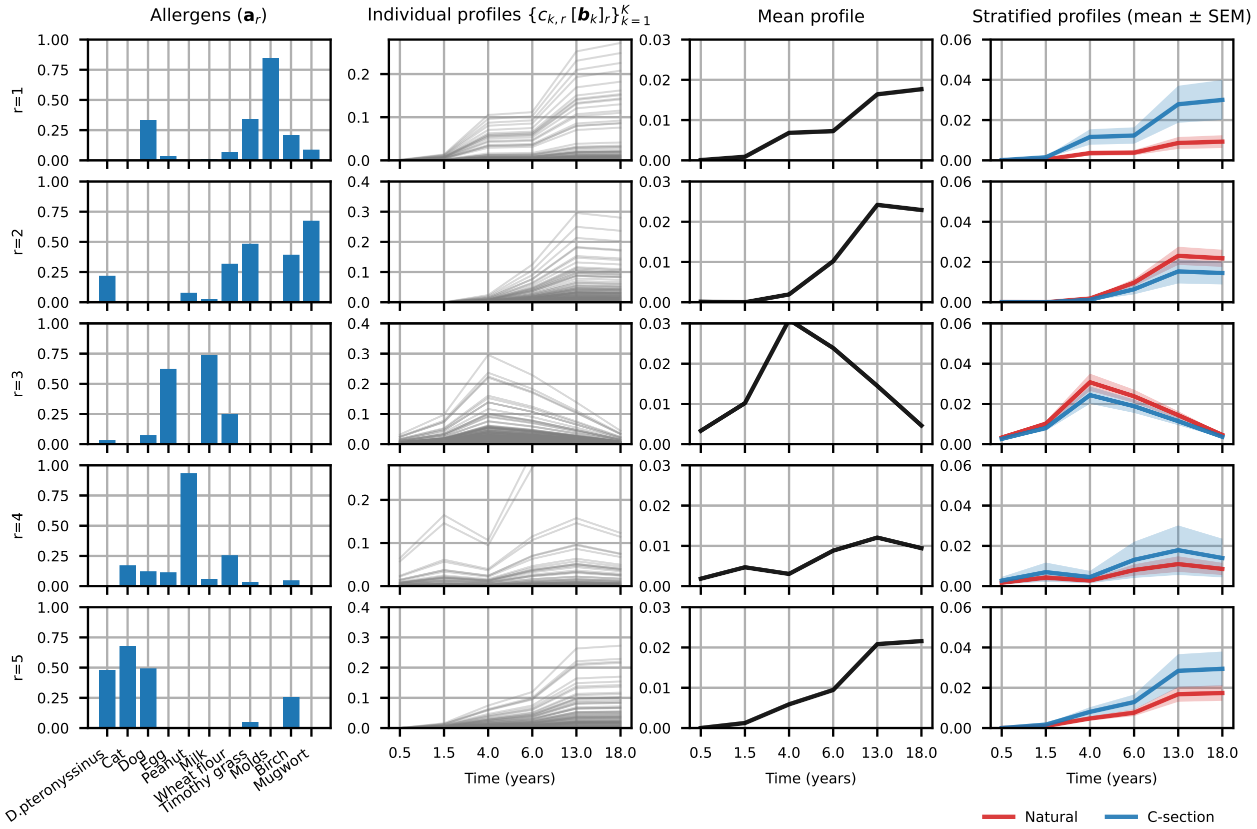


Figure S.9: Sensitization. Components of a 5-component CP model (with nonnegativity constraints in all modes) of the sensitization data. $\mathbf{a}_r$ denotes the pattern in the allergens mode. Subject-specific time profiles scaled by the corresponding subject scores, i.e., $c_{k,r}[\mathbf{b}_k]_r$ are shown in the middle column. Mean of scaled subject-specific profiles are plotted in the third column while the last column shows mean (and standard error of mean) patterns of scaled subject-specific time profiles colored according to delivery/birth mode groups.

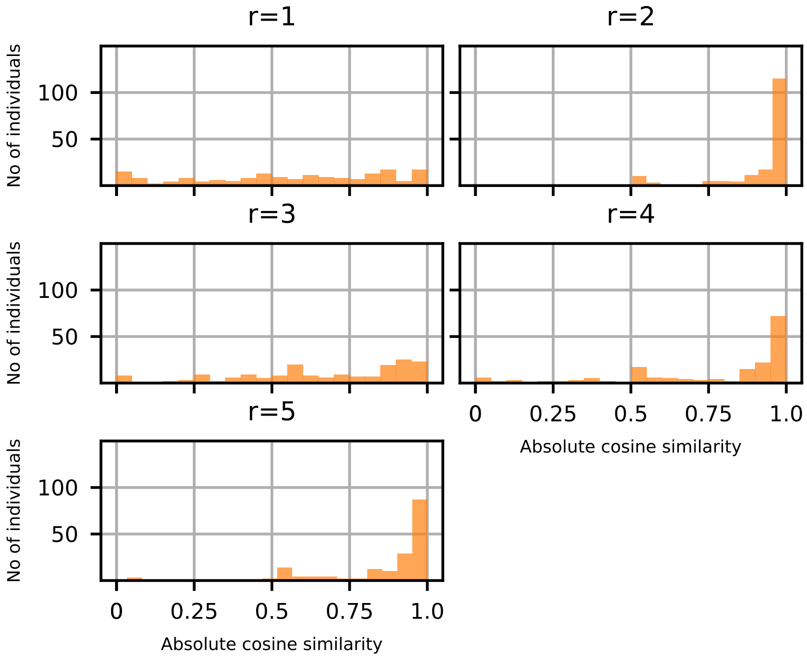


Figure S.10: Comparison of subject-specific time profiles ($c_{k,r}[\mathbf{b}_k]_r$) captured by CMF and PARAFAC2 for each component for sensitization, where each histogram contains 176 data points, one for each subject. The order is adjusted (following the order in the CMF model) so that matching components from different models are compared.
